# Ancient genomes document multiple waves of migration in Southeast Asian prehistory

**DOI:** 10.1101/279646

**Authors:** Mark Lipson, Olivia Cheronet, Swapan Mallick, Nadin Rohland, Marc Oxenham, Michael Pietrusewsky, Thomas Oliver Pryce, Anna Willis, Hirofumi Matsumura, Hallie Buckley, Kate Domett, Nguyen Giang Hai, Trinh Hoang Hiep, Aung Aung Kyaw, Tin Tin Win, Baptiste Pradier, Nasreen Broomandkhoshbacht, Francesca Candilio, Piya Changmai, Daniel Fernandes, Matthew Ferry, Beatriz Gamarra, Eadaoin Harney, Jatupol Kampuansai, Wibhu Kutanan, Megan Michel, Mario Novak, Jonas Oppenheimer, Kendra Sirak, Kristin Stewardson, Zhao Zhang, Pavel Flegontov, Ron Pinhasi, David Reich

**Author notes:** These authors equally supervised this work (Ron Pinhasi, David Reich).

## Abstract

Southeast Asia is home to rich human genetic and linguistic diversity, but the details of past population movements in the region are not well known. Here, we report genome-wide ancient DNA data from thirteen Southeast Asian individuals spanning from the Neolithic period through the Iron Age (4100–1700 years ago). Early agriculturalists from Man Bac in Vietnam possessed a mixture of East Asian (southern Chinese farmer) and deeply diverged eastern Eurasian (hunter-gatherer) ancestry characteristic of Austroasiatic speakers, with similar ancestry as far south as Indonesia providing evidence for an expansive initial spread of Austroasiatic languages. In a striking parallel with Europe, later sites from across the region show closer connections to present-day majority groups, reflecting a second major influx of migrants by the time of the Bronze Age.

---

The archaeological record of Southeast Asia documents a complex history of human occupation, with the first archaic hominins arriving at least 1.6 million years ago (yBP) and anatomically modern humans becoming widely established by 50,000 yBP [*1–3*]. Particularly profound changes in human culture were propelled by the spread of agriculture. Rice farming began in the region approximately 4500–4000 yBP and was accompanied by a relatively uniform and widespread suite of tools and pottery styles showing connections to southern China [*4–6*]. It has been hypothesized that this cultural transition was effected by a migration of people who were not closely related to the indigenous hunter-gatherers of Southeast Asia [*5, 7–10*] and who may have spoken Austroasiatic languages, which today have a wide but fragmented distribution in the region [*4, 5, 11–14*]. In this scenario, the languages spoken by the majority of present-day people in Southeast Asia (e.g., Lao, Thai, Burmese, Malay) reflect later population movements. However, no genetic study has resolved the extent to which the spread of agriculture into the region and subsequent cultural and technological shifts were achieved by movement of people or ideas.

Here we analyze samples from five ancient sites (Table 1; Figure 1A): Man Bac (Vietnam, Neolithic; 4100–3600 yBP), Nui Nap (Vietnam, Bronze Age; 2100–1900 yBP), Oakaie 1 (Myanmar, Late Neolithic/Bronze Age; 3200–2700 yBP [*15*]), Ban Chiang (Thailand, Bronze Age portion of site; 3000–2800 yBP [*16*]), and Vat Komnou (Cambodia, Iron Age; 1900–1700 yBP [*17*]). We initially screened a total of 267 next-generation sequencing libraries from 133 distinct individuals, obtaining powder from petrous bone samples (specifically the high-yield cochlear region [*18*]). For libraries with evidence of authentic ancient DNA, we generated genome-wide data using in-solution enrichment (“1240k SNP” target set [*19*]), yielding sequences from thirteen individuals (Materials and Methods; Table 1; Table S1). Because of poor bone preservation conditions in tropical environments, we observed both a low rate of conversion of screened samples to working data and also limited depth of coverage per sample, and thus we created multiple libraries per individual (66 in total in our final data set).

**Figure 1.**
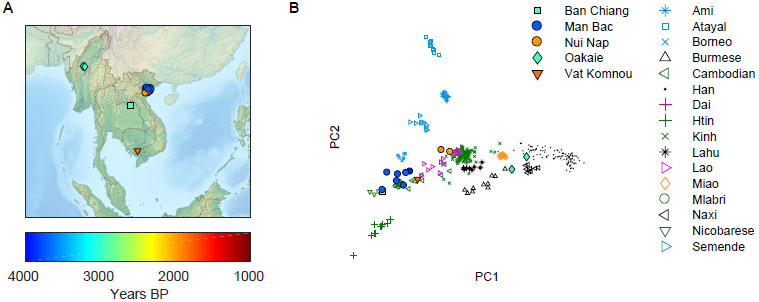
Overview of samples. (A) Locations and dates of ancient individuals. Overlapping positions are shifted slightly for visibility. (B) PCA with East and Southeast Asians. We projected the ancient samples onto axes computed using the present-day populations (with the exception of Mlabri, who were projected instead due to their large population-specific drift). Present-day colors indicate language family affiliation: green, Austroasiatic; blue, Austronesian; orange, Hmong-Mien; black,Sino-Tibetan; magenta, Tai-Kadai.

**Table 1.** Sample information

| ID | Lib. | Date (yBP) | Site | Country/period | Lat. | Long. | Sex | Mt Hap | Y Hap | Cov. |
| --- | --- | --- | --- | --- | --- | --- | --- | --- | --- | --- |
| VN22 | 6 | **3835–3695** | Man Bac | Vietnam NE | 20.1 | 106.0 | F | M13b | .. | 0.048 |
| VN29 | 4 | 3900–3600 | Man Bac | Vietnam NE | 20.1 | 106.0 | F | M7b | .. | 0.022 |
| VN33 | 2 | 3900–3600 | Man Bac | Vietnam NE | 20.1 | 106.0 | M | B5a1a | O | 0.028 |
| VN34 | 10 | **4080–3845** | Man Bac | Vietnam NE | 20.1 | 106.0 | F | M7b1a1 | .. | 0.106 |
| VN37 | 4 | **3825–3635** | Man Bac | Vietnam NE | 20.1 | 106.0 | M | M7b1a1 | CT | 0.019 |
| VN39 | 11 | **3830–3695** | Man Bac | Vietnam NE | 20.1 | 106.0 | M | M7b1a1 | O2a1 | 0.102 |
| VN40 | 6 | **3820–3615** | Man Bac | Vietnam NE | 20.1 | 106.0 | M | M74b | I2a2a2a | 0.041 |
| VN41 | 5 | 2100–1900 | Nui Nap | Vietnam BA | 19.8 | 105.8 | F | C7a | .. | 0.373 |
| VN42 | 7 | **1995–1900** | Nui Nap | Vietnam BA | 19.8 | 105.8 | M | M8a2 | F | 0.055 |
| OAI1/S28 | 5 | 3200–2700 | Oakaie 1 | Myanmar LNBA | 22.4 | 95.0 | F | R | .. | 0.073 |
| OAI1/S29 | 1 | 3200–2700 | Oakaie 1 | Myanmar LNBA | 22.4 | 95.0 | F |  | .. | 0.004 |
| BC8 | 1 | 3000–2800 | Ban Chiang | Thailand BA | 17.4 | 103.2 | F | M | .. | 0.003 |
| AB40 | 4 | **1890–1730** | Vat Komnou | Cambodia IA | 11.0 | 105.0 | M | R | CT | 0.026 |

To obtain a broad-scale overview of the data, we performed principal component analysis (PCA) with a set of diverse non-African populations (East and Southeast Asian, Australasian, Central American, and European [*20–22*]). When projected onto the first two axes, the ancient individuals fall close to present-day Chinese and Vietnamese, with Man Bac shifted slightly in the direction of Onge (Andaman Islanders) and Papuan (Figure S1). Next, we carried out a second PCA using a panel of 16 present-day populations from East and Southeast Asia [*22, 23*]. The populations fill a roughly triangular space in the first two dimensions (Figure 1B; compare [*24*]), with Han Chinese on the right, most Austroasiatic-speaking groups (Mlabri and Htin from Thailand, Nicobarese, and Cambodian, but not Kinh) toward the left, and aboriginal (Austronesian-speaking) Taiwanese at the top. Man Bac, Ban Chiang, and Vat Komnou cluster with Austroasiatic speakers,while Nui Nap projects close to present-day Vietnamese and Dai near the center, and Oakaie projects close to present-day Burmese and other Sino-Tibetan speakers (two samples, BC8 and OAI1/S29, have especially low coverage, so their exact positions should be interpreted with caution). Present-day Lao are intermediate between Austroasiatic speakers and Dai, and the Borneo and Semende populations from western Indonesia fall intermediate between Austroasiatic speakers and aboriginal Taiwanese.

We measured levels of allele sharing between populations via outgroup *f_3_*-statistics and observed results consistent with those from PCA (Table S2). Nominally, the top sharing for each ancient population is provided by another ancient population, but this pattern is likely to be an artifact due to correlated genotype biases between different ancient samples (Supplementary Text). Restricting to present-day comparisons, Man Bac shares the most alleles with Austroasiatic-speaking groups (as Austroasiatic-speaking groups do with each other), Nui Nap with Austronesian speakers and Dai, Oakaie with Sino-Tibetan-speaking groups, and Vat Komnou with a range of different populations. We also investigated the relationships between the ancient individuals and archaic hominins (Neanderthal and Denisova). Using Han Chinese as a baseline, we observed nominal signals of excess archaic alleles in all populations (statistically significant for Man Bac and Nui Nap; all 1240k SNPs), but as above, these results appear to be driven by artifacts in the data, rather than reflecting actual excess archaic ancestry (Supplementary Text).

The genetic clustering of the early farmer samples with Austroasiatic-speaking populations could be due to shared genetic drift along a common ancestral lineage, shared ancestral admixture, or both. We computed statistics of the form *f_4_* (*X, SEA*; *AUS,NEA*)—with *SEA, AUS*, and *NEA* being reference Southeast Asian, Australasian (typically a union of Papuan and Onge), and Northeast Asian populations, respectively—which take increasingly positive values for increasing proportions of deeply-splitting ancestry (from outside the East Asian clade) in test population *X*. Figure 2 shows values of *f_4_* (*X*, Kinh; Australasian, Han) for present-day and ancient populations. Isolated Austroasiatic speakers yield significantly positive statistics, as do the majority of the ancient samples, with approximately equal values for Mlabri, Nicobarese, and Man Bac. The Man Bac individuals are additionally mostly similar to each other, except for one, VN29, which is significantly higher than the population mean (Bonferroni-corrected *p* = 0.025), with VN40 also modestly higher (overall homogeneity rejected by χ ^2^ = 15.8, *p <* 0.02; Materials and Methods). Vat Komnou and Oakaie OAI1/S28 are also positive, as are present-day Cambodian and Burmese, while Nui Nap is close to zero (*Z* = 1.2).

**Figure 2.**
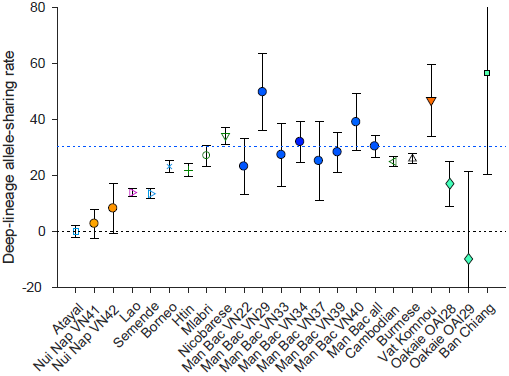
Relative amounts of deeply diverged ancestry. The Y-axis shows *f_4_* (*X*, Kinh;Australasian, Han) (multiplied by 10^4^) for populations *X* listed on the X-axis (present-day as aggregate; ancient samples individually, except for “Man Bac all”).Symbols are as in Figure 1. Bars give two standard errors in each direction; dotted lines indicated the levels in Man Bac (top, blue) and Kinh (zero, black).

Using these observations as a starting point, we built admixture graph models to test the relationships between the Vietnam Neolithic samples and present-day Southeast Asians in a phylogenetic framework. We began with a scaffold model containing the Upper Paleolithic Siberian Ust’-Ishim individual as an outgroup and present-day Mixe, Onge, and Atayal. We then added Nicobarese and Mlabri, two present-day Austroasiatic-speaking populations that appear to have relatively simple admixture histories, as well as Man Bac. All three are inferred to have ancestry from a Southeast Asian farmer-related source (∼70%, forming a clade with Atayal) and a deeply diverging eastern Eurasian source (∼30%, sharing a small amount of drift with Onge; *f*-statistics indicate that this source is also not closely related to Papuan or South Asians). The allele sharing demonstrated by outgroup *f_3_*-statistics can be explained in the admixture graphs by shared genetic drift along the farmer lineage, along the deeply-splitting lineage, or both, but we are not able to determine the relative contributions without additional unadmixed reference populations (Supplementary Text). Given the closeness of the mixture proportions among the three groups, however, we found that the most parsimonious model (Figure 3; Figure S2) involved a shared ancestral admixture event (29% deep ancestry; 28% omitting VN29), followed by divergence of Man Bac from the present-day Austroasiatic speakers, and finally a second pulse of deep ancestry (5%) into Nicobarese. We did not fit full models for the more recent samples given their thinner coverage and likely more complex histories; however, we used *f*-statistics to probe the relationship between Nui Nap and present-day Southeast Asians more carefully. We found that the statistic *f_4_* (Nui Nap, *X*;*Y, Z*) is slightly but significantly different from 0 (|*Z|* > 2.3) for any combination of 1000Genomes East Asian populations *X, Y*, and *Z* (Dai, Kinh, Han, or Japanese; all 1240k SNPs), pointing to small but measurable changes in ancestry between the Bronze Age period in Vietnam and today.

**Figure 3.**
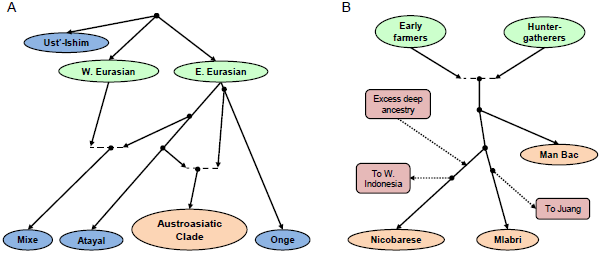
Schematics of admixture graph results. (A) Wider phylogenetic context. (B) Details of the Austroasiatic clade. Branch lengths are not to scale, and the order of the two events on the Nicobarese lineage in (B) is not well determined (Supplementary Text).

Finally, to shed more light on early divergences of geographically diverse Austroasiatic lineages, we fit an extended admixture graph with additional populations. First, while western Indonesians (here represented by Borneo and Semende) speak Austronesian languages, they appear, as mentioned above and as previously hypothesized [*25, 26*], to be admixed with ancestry derived from both Austronesian- and Austroasiatic-associated sources. We confirmed that both Borneo and Semende fit well in the admixture graph with these two components, although a small proportion of a third, deeply-splitting (likely indigenous) component was necessary as well (Figure S3; Supplementary Text). Borneo was inferred to have ∼38%, 59% and 3% of the Austronesian, Austroasiatic, and indigenous components, respectively, while Semende was inferred to have ∼67%, 29% and 4%, with the Austroasiatic component closer to Nicobarese than to Mlabri or Man Bac, forming a “southern” Austroasiatic sub-clade (Figure 3B). Second, for Juang, an Austroasiatic-speaking population from India [*22*], we also obtained a good fit with three ancestry components: one western Eurasian, one deep eastern Eurasian (interpreted as an indigenous South Asian lineage), and one from the Austroasiatic clade (Figure S3). The Austroasiatic source for Juang (proportion 37%) is inferred to be closest to Mlabri, as supported by statistics *f_4_* (Juang, Palliyar; Mlabri, *X*) > 0 for *X* = Atayal, Man Bac, or Nicobarese (*Z* = 5.1, 2.8, 2.3), creating a “northern” sub-clade.

Our results provide strong genetic support for the hypothesis that agriculture was first practiced in Mainland Southeast Asia by (proto-) Austroasiatic-speaking migrants from southern China [*4–6, 11–13*]. We find that all seven of our sampled individuals from Man Bac are closely related to present-day Austroasiatic speakers, including a shared pattern of admixture, with one, VN29, showing significantly elevated indigenous ancestry. By comparison, studies of cranial and dental morphology have placed Man Bac either close to present-day East and Southeast Asians (“Neolithic”), intermediate between East Asians and a cluster containing more ancient hunter-gatherers from the region plus present-day Onge and Papuan (“indigenous”), or split between the two clusters [*7,9,27*]. The simplest explanation for our results is that the majority of our samples represent a homogeneous Neolithic cluster, with recent local contact between farmers and hunter-gatherers leading to additional hunter-gatherer ancestry in VN29 and perhaps VN40 [*7, 9*]. This model would imply that the incoming farmers had already acquired 25–30% hunter-gatherer ancestry, either in China or Southeast Asia, establishing the characteristic Austroasiatic affiliated genetic profile seen in multiple populations today. The symmetric position with respect to Native Americans of (1) the majority East Asian ancestral lineage in Man Bac, and (2) aboriginal Taiwanese, points to an origin for the farming migration specifically in southern China (contrasting with *f_4_* (*X*, Atayal; Mixe, Dinka) > 0 for northern East Asians *X* = Han, Japanese, or Korean, *Z* > 4.5).

Our findings also have implications for genetic transformations linked to later cultural and linguistic shifts in Southeast Asia and beyond. We observe substantial genetic turnover between the Neolithic period and Bronze Age in Vietnam, likely reflecting a new influx of migrants from China [*28*], although it is striking that present-day majority Vietnamese (Kinh), who are closely related to our Bronze Age samples from Nui Nap, still speak an Austroasiatic language. Late Neolithic/Bronze Age individuals from Oakaie also do not possess an Austroasiatic genetic signature, in their case being closer to populations speaking Sino-Tibetan languages (including present-day Burmese), pointing to an independent East Asian origin. Outside of Mainland Southeast Asia, we document admixture events involving derived Austroasiatic-related lineages in India (where Austroasiatic languages continue to be spoken) and western Indonesia (where all languages today are Austronesian), with the link between Borneo, Sumatra and Nicobarese supporting archaeological hints of an initial Austroasiatic-associated Neolithic settlement of western Indonesia [*26*]. Overall, Southeast Asia shares common themes with Europe, Oceania, and sub-Saharan Africa, where ancient DNA studies of farming expansions and language shifts have revealed similarly high levels of genetic turnover associated with archaeologically attested transitions in culture.

## Acknowledgments

We thank Iosif Lazaridis, Vagheesh Narasimhan, Iñigo Olalde, and Nick Patterson for technical assistance; Nicole Adamski and Ann-Marie Lawson for aiding with lab work; and Minh Tran Thi, Rona Ikehara-Quebral, Miriam Stark, Michele Toomay Douglas, and Joyce White for help with archaeological samples. **Funding:** This work was supported by the French Ministry for Europe and Foreign Affairs (T.O.P.), National Science Foundation (HOMINID grant BCS-1032255; D.R.), National Institutes of Health (NIGMS grant GM100233; D.R.), an Allen Discovery Center of the Paul Allen Foundation (D.R.), and the Howard Hughes Medical Institute (D.R.). **Author contributions:** N.R., P.F., R.P., and D.R. supervised the study. M.O., M.P., T.O.P., A.W., H.M., H.B., K.D., N.G.H., T.H.H., A.A.K., T.T.W., B.P., and R.P. provided samples and assembled archaeological and anthropological information. M.L., O.C., S.M., N.R., N.B., F.C., D.F., M.F., B.G.,E.H., M.M., M.N., J.O., K.Si., K.St., Z.Z., R.P., and D.R. performed ancient DNA laboratory and data processing work. P.C., J.K., W.K., and P.F. provided present-day data. M.L., S.M., and D.R. analyzed genetic data. M.L., R.P., and D.R. wrote the manuscript with input from all coauthors. **Competing interests:** The authors declare no competing interests. **Data and materials availability:** The aligned sequences will be available through the European Nucleotide Archive under accession number PRJEB24939. Genotype datasets used in analysis will be available at https://reich.hms.harvard.edu/datasets.

## Materials and Methods

### Experimental design

DNA was extracted from archaeological samples as described below. Resulting genotype data were analyzed using population genetic tools to infer historical processes.

### Ancient sample preparation and data processing

We screened a total of 133 ancient petrous bone samples for the presence of human DNA, following an established procedure [*19, 29, 30*]. We obtained bone powder in a dedicated clean room facility at University College Dublin and extracted DNA via published protocols [*31, 32*] in clean rooms at Harvard Medical School. A subset of the extracts were executed using silica magnetic beads instead of the standard silica spin columns (Table S1). From the extracts, we prepared double-stranded individually bar-coded libraries, some of which (including all libraries used for final analyses aside from Ban Chiang) we treated with uracil-DNA glycosylase (partial UDG treatment) to reduce the rate of characteristic cytosine-to-thymine errors in ancient DNA [*33, 34*]. For the majority of libraries, we used magnetic bead cleanup between enzymatic reactions and SPRI bead cleanup for the final PCR [*35, 36*] instead of MinElute column cleanups (Table S1). We initially used target capture hybridization to enrich the libraries for sequences overlapping the mitochondrial genome [*37, 38*] and in most cases a set of approximately 3000 nuclear SNP targets, and we then sequenced the enriched libraries on an Illumina NextSeq 500 instrument with 76-base-pair paired-end reads. From the output, we merged sequences that were within 1-base-pair edit distance of expected bar-codes and with at least 15 overlapping bases, trimmed bar-codes and adapters, and mapped the merged reads to the mitochondrial reference genome RSRS [*39*] or to the human reference genome (version hg19) as appropriate. Mapped reads were then quality-filtered and de-duped, and two terminal bases were clipped to reduce damage (five for UDG-minus libraries), as described previously [*29*]. For libraries with evidence of nuclear DNA, we then enriched for sequences overlapping approximately 1.2 million genome-wide SNPs [*19, 30, 40*] (in some cases pooling libraries from the same individual prior to enrichment; Table S1) and sequenced to increased depth, processing the data in the same way. We called one allele at random per site to create pseudo-haploid genotypes and determined genetic sex by examining the factions of reads mapping to the X and Y chromosomes.

Because of the poor molecular preservation of the samples, we prepared multiple libraries for most individuals (66 libraries used in final analyses for 13 samples, out of 128 libraries screened for those samples; Table S1), which we then merged after data processing. All 66 libraries displayed ancient DNA damage (at least 16% C-to-T substitutions in the final base of mitochondrial screening sequencing reads), providing evidence of authenticity [*34, 41*], with noticeably high damage rates for these samples likely reflecting hot and humid local climates (Supplementary Text). During screening, we assessed possible contamination by measuring rates of apparent heterozygosity on the mitochondrial genome [*40*], and we performed follow-up heuristic analyses on the genome-wide data to test for the presence of potentially contaminating present-day human DNA (Supplementary Text).

Mitochondrial DNA haplogroups were called with HaploGrep2 [*42*] using phylotree(mtDNA tree Build 17; 18 Feb 2016), with final calls based on comparisons between single-library results and reassembled multi-library merges for each individual (versions with all reads and with only reads showing evidence of damage). Y-chromosome haplogroups were determined from 15,100 targeted SNPs; mutations were compared with the tree provided by the International Society of Genetic Genealogy (http://www.isogg.org) via a modifiedversion of the yHaplo software [*43*].

### Present-day data

We generated new genome-wide SNP genotype data for 10 Htin and 10 Mlabri individuals [*24, 44, 45*] (who previously gave informed consent for genome-wide analyses of population history and public sharing of anonymized data following publication) on the Human Origins Array. We merged these new data with published Human Origins samples [*20, 22, 23, 46, 47*] and with 1000 Genomes populations [*21*]. Han, Kinh, and Japanese data were taken from 1000 Genomes, whereas Dai were taken from Human Origins, except for the statistic *f_4_* (Nui Nap, *X*; *Y, Z*), where we used all 1000 Genomes populations to increase power and retain symmetry of data sources. All analyses were performed using the set of 593,124 autosomal Human Origins SNPs, unless otherwise noted (“all 1240k SNPs” refers to the full set of about 1.15 million targeted autosomal SNPs).

### Statistical analysis

We performed PCA by computing principal components for present-day populations (except as noted) and then projecting ancient samples, using the “lsqproject” and “autoshrink” options in smartpca [*48, 49*]. Admixture graphs and *f*-statistics were implemented via ADMIXTOOLS [*20*] (differences between *f*-statistics using the qp4diff program with “allsnps” mode), with standard errors estimated via block jackknife. To test for homogeneity of *f*-statistics, we computed the sum of the squares of the *Z*-scores of differences between each individual in a population and the aggregate population value,which has a *χ ^2^n-*1distribution under the null, as in a *χ ^2^* test for variance (both heterogeneity statistics for VN29 were also replicated at *p <* 0.05 using full sequence data for Papuan and Andamanese [*50, 51*] and all 1240k SNPs). We note that no individuals in the study were identified as close relatives based on allele matching rates; for the coverage level of the Man Bac samples, we can confidently rule out any first-degree kinship but not more distant relationships.

## Supplementary Text

### Analysis of possible contamination

Our initial methods to authenticate our data and estimate levels of possible contamination were based on established protocols implemented in our screening process. First, we observed characteristic ancient DNA damage patterns, with at least 16% C-to-T substitutions in terminal positions of molecules mapping to mtDNA (roughly twice the rate in non-UDG-treated libraries versus partial UDG libraries: min 37%, mean 62%, median 62%, max 74% for all 29 non-UDG libraries; mean 32%, median 33%, max 43% for 65 partial-UDG libraries used in analyses). Such high damage rates make it less likely that the samples have substantial amounts of contamination (especially for the important VN29 sample, with damage rates of ∼40% in all four libraries) but do not provide quantitative estimates. We also measured apparent heterozygosity at single-copy markers (mtDNA as well as the X chromosome in males), with mtDNA results shown in Table S1. Of the 66 libraries used in analyses, 39 yielded estimates of the mismatch rate, with a range of 0.1–49.5% (0.1–20.5% excluding one outlier library, mean 7.2%, median 5.4%). However, based on previous experience, we believe that the exact quantitative results are not always reliable, especially for low-coverage libraries. Estimates based on the X chromosome are generally more stable, but we lacked sufficient coverage for the samples in this study to generate confident measurements with this method.

While the damage patterns and mtDNA matching results indicated reasonably good quality data, we wished to extend our quality control by examining potential effects of contaminating DNA on observed population genetic results. First, we evaluated the positions of the ancient samples in PCA in light of possible contamination. In the broad scale PCA (Figure S1), most samples are very close to present-day East and Southeast Asians, with only AB40 (Vat Komnou, Cambodia Iron Age) perhaps showing signs of greater affinity to Europeans, which could be a result of a small amount of contamination (or could reflect actual western Eurasian ancestry, conceivably via India). It is also possible that contamination could come from a different source, with the other most likely ancestry being East Asian, which would be difficult to detect in Figure S1. Thus, we turned next to our PCA focusing on East and Southeast Asia (Figure 1B). Here, any expected effects would be more subtle, but none of the samples appear to be shifted unexpectedly in the direction of present-day populations such as Han or Kinh, and the ancient populations are all relatively homogeneous. We also projected versions of the sample data restricted to sequencing reads showing ancient DNA damage patterns, but we found that the coverage was too low (generally at least 10 times thinner than the already low-coverage full data) to draw any informative conclusions from PCA (or other analyses).

Finally, we computed *f*-statistics designed to detect excess affinity to potentially contaminating populations. First, the statistic *f_4_* (*X*, Kinh; European, Yoruba) is close to zero (*|Z| <* 2.1) for all samples *X*. A weak signal of European contamination could be masked by affinity to the African outgroup (see next section), but we do not believe that any of the samples could be heavily affected. Next, we computed statistics *f_4_* (*X, Y*; *Z, W*), where *Y, Z, W* are any permutation of Han, Japanese, and Kinh (the most likely potential East/Southeast Asian contamination sources). These patterns are more complicated to interpret, but we believe they support the authenticity of the data. Here we were most interested in the results for the Man Bac samples, which we expect to have relatively uniform affinity to the present-day East and Southeast Asians. Indeed, the seven individuals yield values of *f_4_* (*X*, Han; Kinh, Japanese) = 0.0025–0.0034, *f_4_* (*X*, Japanese; Kinh, Han) = 0.0025–0.0031, and *f_4_* (*X*, Kinh; Han, Japanese) = 0–0.0005, with standard errors of 0.0001–0.00025. In comparison to the observed results, if we substitute the present-day populations themselves in the first position, we find that *f_4_* (Kinh, Han;Kinh, Japanese) = 0.0076, *f_4_* (Han, Japanese; Kinh, Han) = 0.0009, and *f_4_* (Japanese,Kinh; Han, Japanese) = *-*0.0087, plus *f_4_* (*Y, Y*; *Z, W*) = 0 for any *Y*. The differences between these sets of values imply that any substantial genetic material introduced from one of the present-day populations would be noticeable in the sample(s) affected. A trace amount of contamination could be present, but (by definition) this would not significantly affect our results. We note in particular that if the VN29 sample were contaminated with present-day East Asian DNA, this would cause its apparent proportion of deeply splitting ancestry to be too low, preserving our observation of within-site heterogeneity.

### Signals related to potential data artifacts

We observed two phenomena that we believe to be due to data artifacts: (1) excess affinity between different ancient samples, and (2) excess affinity between ancient samples and deep outgroups. Observation (1) is reflected in the fact that every ancient population apparently shares the most alleles with another ancient population (via outgroup *f_3_*statistics), even when the pair do not belong to the same genetic cluster. For example, we find *f_3_* (Dinka; Man Bac, Nui Nap) = 0.206, *f_3_* (Dinka; Man Bac, *AA*) = 0.189 (where *AA* refers to Mlabri, Nicobarese, or Htin, who have the greatest allele sharing with Man Bac among present-day populations), and *f_3_* (Dinka; Man Bac, *EA*) *≥* 0.173 for any other East/Southeast Asian population *EA*. Such a large gap between Nui Nap and all present-day populations, including some that are closely related to Nui Nap (such as Kinh), seems highly implausible. For observation (2), we found, for example, that all statistics *f_4_* (*X*, Han; Altai/Denisova, Yoruba) are positive for ancient populations *X*, reaching statistical significance for Man Bac and Nui Nap (*Z >* 4 and *Z >* 2.5, respectively; all 1240k SNPs). However, both populations display even greater allele sharing with chimpanzee (*f_4_* (Man Bac/Nui Nap, Han; Altai/Denisova, Chimp) *<* 0), which would not be expected if they harbored Neanderthal or Denisova-related ancestry; by comparison, the statistics *f_4_* (Papuan, Han; Altai/Denisova, Chimp) are strongly positive (*Z >* 12 for Denisova and *Z >* 4 for Altai Neanderthal). While we cannot formally rule out a component of unknown archaic ancestry, we believe that these signals are again influenced by artifacts in the data, in this case leading to excess ancestral allele calls. As a result of such potential artifacts, we attempt to minimize the use of deep outgroups or unbalanced comparisons between ancient samples. (We note that for outgroup *f_3_*-statistics, we used either Dinka or Europeans (CEU) as the outgroup and obtained concordant results.)

## Details of admixture graph fitting

### Core model

To simplify the fitting of Mixe, we locked its western Eurasian ancestry proportion at 30%. We also obtained similar results when replacing Mixe with Ulchi from the Amur River Basin (∼5% western Eurasian ancestry).

Without additional unadmixed reference populations available in our admixture graphs, we did not have power to resolve the exact topology of the two source lineages for Austroasiatic-clade populations. The final model we present, with an initial shared ad mixture event for Nicobarese, Mlabri, and Man Bac, is the most parsimonious version, but others are possible as well. In particular, we can also fit all three populations with separate admixture events, which yields a very similar fit score but with two additional free parameters in the model. In these less-parsimonious models, we cannot distinguish between versions in which (1) the farmer ancestry source is the same for all three and the deep ancestry sources are different, or (2) the farmer ancestry sources are diverged and the deep ancestry sources are the same for all three. We note though that in all of the different models, Man Bac is inferred to diverge prior to the split of Nicobarese from Mlabri. For our final topology, we also tested the possibility that Man Bac and Nicobarese could share the same mixture proportion, with a small additional pulse of farmer ancestry into Mlabri rather than a pulse of deep ancestry into Nicobarese, and while it was not strongly rejected, this version had a fit score (approximate log-likelihood) roughly 3 worse, with a residual statistic at *Z* = 2.0 indicating a difference in ancestry proportions between Man Bac and Nicobarese.

Several other lines of evidence also led us to prefer the model with an initial shared admixture event for the Austroasiatic clade. As discussed in the main text, the structure of our population sample from Man Bac is suggestive of a group of immigrant farmers who had already experienced admixture with hunter-gatherers. It is also notable that Nicobarese, Mlabri, and Man Bac have such similar mixture proportions despite their wide geographic separation. There are reasons moreover to think that even if these three groups are descended (partially or completely) from separate farmer/hunter-gatherer ad mixture events, the farmer lineages represent a single migration out of China (most likely associated with Austroasiatic languages). Formally, we cannot prove this assertion, as we did find one less-parsimonious model (with statistically indistinguishable fit quality) in which the farmer component for Man Bac can be fit outside the Austroasiatic clade (or even the Austroasiatic-plus-Austronesian clade). However, the most likely alternative source of farmer ancestry for Man Bac would be the Austronesian expansion, which is associated with a strong genetic drift signal that we would almost certainly be able to detect (as in the observed differences between Man Bac and Austroasiatic speakers versus western Indonesians in PCA and other analyses). We also used *f*-statistics to assess the relationships of Nicobarese, Mlabri, and Man Bac with respect to other present-day nonAustroasiatic-speaking populations from Southeast Asia, and we found no evidence of asymmetry. Finally, given that the deep ancestry in Austroasiatic speakers is not closely related to Onge, Papuan, or other Australasians, it seems more likely that this ancestry would have been acquired in a coordinated way throughout the clade rather than requiring separate admixture events that nonetheless involved very similar sources of deep ancestry for Austroasiatic speakers in different parts of Southeast Asia.

Lastly, we note that we fit a model for Man Bac with all 1240k SNPs, using full sequence data for other present-day populations, but with no other Austroasiatic speakers present, and the results (insofar as the models were overlapping) were very similar.

### VN29

Because of the small number of SNPs covered, we could not model VN29 and the remainder of the Man Bac individuals simultaneously. We did fit a version with VN29 alone in place of Man Bac, this time using a model with separate admixture events, and as expected, it was inferred to have more deep-lineage ancestry than Nicobarese and Mlabri, with a point estimate of 37%. The topology was identical to that for all of Man Bac together, although some of the parameters (branch lengths and mixture proportions) differed modestly from the core model, given the different set of SNPs.

### Western Indonesians

To help resolve the different sources of ancestry in western Indonesians, we also added Dai to our admixture graph model. Dai are modeled as three-way admixed (evidence of admixture provided by negative *f_3_* statistics, e.g., *f_3_* (Dai; Han, Nicobarese) = *-*0.0014, *Z* = *-*3.3), with the majority component closely related to the farmer ancestry in Austroasiatic speakers, plus small proportions of northern East Asian (10%) and deep eastern Eurasian (10%, same source as in Austroasiatic speakers) ancestry. We do not necessarily believe that these represent proximal mixing populations, but the resulting fit was reasonable and satisfactory, and the exact (likely complex) history for Dai is not directly relevant for our study.

For Borneo and Semende, we tested a number of alternative models in addition to the final version presented. Using our main graph topology, we obtained a significantly better fit for Borneo as a mixture of Austroasiatic-clade and Austronesian (plus additional indigenous) ancestry as compared to only Austronesian and indigenous, even if the deeply-splitting ancestry is the same that contributes to Austroasiatic speakers, and also allowing for a combination two different deep ancestry components. In a less-parsimonious topology for the Austroasiatic clade, with separate deep ancestry sources for Nicobarese, Mlabri, Man Bac, and western Indonesians, a simpler Austronesian-plus-indigenous model does fit successfully for Borneo or Semende individually, but it is again worse when modeling Borneo and Semende together. We also note that in the final version, we modeled the Austroasiatic-related (Nicobarese-related) component in western Indonesians as including the extra deep indigenous ancestry present in Nicobarese, but the alternative model (western Indonesians having ancestry splitting from the Nicobarese lineage prior to the second admixture event) also fits well, so we do not have the resolution to determine the order of those two events.

### Juang

For Juang, the inferred western Eurasian component is almost certainly itself admixed, but for the purposes of our model, it fits best as closely related to the Ancient Northern Eurasian lineage forming part of the ancestry of Native Americans. The deep eastern Eurasian component splits close to the same point as Onge, East Asians, and the indigenous Austroasiatic component. A mixture of two components of this type is characteristic of Indian populations today; Juang, however, also traces ancestry to a third, Austroasiatic-related source.

## Summary of Archaeologial Context for Sampled Sites

### Man Bac (Vietnam Neolithic)

Man Bac is on the southern edge of the Red River Delta, 25 km from the current coast, in Yen Mo District, Ninh Binh Province, northern Vietnam. The site is well sheltered by surrounding steep karstic outcrops. Man Bac was comprehensively excavated in 1999, 2001, 2004-5 and 2007 revealing a wealth of material cultural and zooarchaeological remains in addition to 100 human burials (Oxenham et al. 2011). While the pottery styles, including manufacture techniques and decoration, have a strong local influence, it is clearly associated with the broadly distributed Phung Nguyen culture in northern Vietnam. While the Phung Nguyen is often identified with the earliest introduction of bronze into northern Vietnam, there is no evidence for a knowledge of bronze at Man Bac, and indeed the dating of the introduction of Bronze into northern Vietnam is simply not known (Oxenham 2015).

The site, while contextually complex, essentially consists of three major stratigraphic units, the upper two being associated with structures (as evidenced by extensive evidence for post holes) and everyday living (hearths, food remains, and general debitage). The lower layer, extending to approximately 2m in depth in parts, is for the most part free of general midden material and otherwise sterile except for the burials. In general, Man Bac displays evidence for elevated levels of fertility, cranio-dental morphological and mtDNA diversity, rice cultivation and domestic pig rearing as well as a broad and diverse continued reliance on hunting as part of the subsistence mix. Mortuary studies of Man Bac have indicated a loose age-based hierarchy and complex system of social identities, including a range of age-based transitions in childhood (Oxenham et al. 2008). Further, Man Bac provided the backdrop to the development of the new subdiscipline “the bioarchaeology of care” with the adult quadriplegic case of Man Bac 09 (Oxenham et al.2009; Tilley and Oxenham 2011).

## Samples used in this study

- VN22 (07.MB.H2.M15): Female, 3836-3694 cal yBP (3490±25 BP, PSUAMS-1920)
- VN29 (05.MB.M16): Female, estimated 3900-3600 yBP
- VN33 (07.MB.H1.M10): Male, estimated 3900-3600 yBP
- VN34 (07.MB.H1.M6): Female, 4080-3845 cal yBP (3630±35 BP, Poz-81116, date suspect due to C:N ratio of 3.68 – but reasonable based on archaeological context, and only slightly elevated C:N ratio)
- VN37 (07.MB.H1.M09): Male, 3825-3637 cal yBP (3445±20 BP, PSUAMS-2409)
- VN39 (07.MB.H2.M16): Male, 3831-3694 cal yBP (3480±20 BP, PSUAMS-2410)
- VN40 (07.MB.H2.M14): Male, 3818-3614 cal yBP (3430±20 BP, PSUAMS-2370)

### Nui Nap (Vietnam Bronze Age)

First surveyed in 1962 and subsequently excavated in 1976-77, Nui Nap is located at the base of a limestone mountain in Dong Hieu District, Thanh Hoa province, only 10km distant from the eponymous Dong Son site. Over 30 extended supine burials were recovered, including a broad range of mortuary offerings, including: bronze spear and arrow heads, daggers, axes, harpoons, vessels, earrings, drums, beads (including glass), pottery and even the occasional Han coin (Oxenham 2016: 10). Nui Nap provides the first verified evidence of the use of betel nut (*Areca catechu)* in northern Vietnam (Oxenham et al. 2002) and along with other Bronze and Iron Age sites in the region, contributes to our understanding of the rise of infectious disease with the emergence of agricultural dependence (Oxenham et al. 2005).

Nui Nap is situated in a region conquered by the Han in the late first millennium BC, becoming the southern-most Han administrative region for much of the first millennium AD. Indeed, there are clear records for massive Han migration into northern Vietnam from the first century AD (Oxenham 2016).

## Samples used in this study

- VN41 (78.NN.M4.KB): Female, estimated 2100-1900 yBP
- VN42 (77.NN.M7.KB): Male, 1994-1901 cal yBP (2005±15 BP, PSUAMS-2371)

### Ban Chiang (Thailand Bronze Age)

Ban Chiang, an UNESCO world heritage archaeological site that spans the pre-metal (Neolithic) to Bronze/Iron Ages, is located in the village of Ban Chiang, Nong Han District, Udon Thani Province, northeastern Thailand (Pietrusewsky & Douglas 2002). Radiocarbon dating, based primarily on artifacts from this site, suggests dates of ca. 2100 BCE to 200 CE (White & Hamilton 2009). Dating of the human and associated animal bones from the site suggests the initial settlement of Ban Chiang occurred ca. 1600-1450 BCE, with the transition to the Bronze Age occurring ca. 1100 BCE (Higham et al. 2015). The archaeological sequence at Ban Chiang is known for its distinctive decorative pottery, ornaments, elaborate burial offerings, and early evidence of metallurgy and agriculture, including bronze artifacts and domesticated rice. A total of 142 burials from two separate sites in the village of Ban Chiang, about 100 meters apart, which were excavated under the direction of Chester Gorman (University of Pennsylvania) and Pisit Charoenwongsa (Thai Fine Arts Department-FAD) in 1974 and 1975, are described in Pietrusewsky and Douglas (2002).

## Sample used in this study

- BC8: Female, estimated 3000-2800 yBP

### Oakaie (Myanmar Late Neolithic/Early Bronze Age)

Partial excavation of the Oakaie 1 (OAI1) cemetery in 2014-15 revealed forty single and six double burials of adults and juveniles, male and female, cut into a sterile volcanic tuff at varying depths and orientations. Funerary offerings included bivalve shells, pottery, stone beads and bracelets, bone bracelets, spindle whorls, a cowrie shell and a dog. Metal was found in only one grave, S15, in the form of a single bronze axe. A complex stratigraphy with significant ancient and recent disturbance currently precludes the definitive attribution of the burials without metal,including S28 and S29, to either the Neolithic or Bronze Age periods. In the absence of preserved collagen, radiocarbon dating was attempted using bone and tooth apatite and shells but most of the determinations are problematic, typically appearing too young^1^. More reliable dating is provided by the extrapolation of charcoal dates and ceramic techno-typologies from closely neighbouring sites.

Indeed, OAI1 is but is one activity area of an extensive late Neolithic to early Bronze Age site located on the eastern bank of the Chindwin, approximately 100 km north of the confluence with the Irrawaddy. Nyaung’gan, the first late prehistoric site investigated by Myanmar archaeologists^2^, lies 2.6 km to the NNE and a vast settlement and industrial zone extends over at least 1000 m to the south, west and north, with excavated locations OAI2-4. A series of 52 ^14^C dates indicate occupation from the 12^th^ to 8^th^ centuries BC, with the Bronze Age transition probably falling in the 10^th^ century^1^. A major lithics industry is evident in the production of axe/adzes, beads, bracelets and other ornaments, partly derived from proximity to sources of volcanic rock but also using imported agate, carnelian, nephrite and other minerals.

Archaeometallurgical analyses indicate onsite secondary production activity (founding) but that the copper used is consistent with imports from central Laos and central Thailand, and not with the nearby deposits at Monywa^3^. These combined data suggest that the population buried at OAI1 had some degree of interaction with groups over in excess of 1000 km of MSEA territory.

## Samples used in this study

- OAI1/S28: Female, estimated 3200-2700 yBP
- OAI1/S29: Female, estimated 3200-2700 yBP

### Vat Komnou (Cambodia Iron Age)

The Vat Komnou cemetery, ca. 200 BCE – 200 CE, at the Angkor Borei site in southern Cambodia is located on the western edge of the Mekong Delta (Ikehara-Quebral et al. 2017; Stark 2006). The dates for this site fall within the Protohistoric Period or Iron Age (ca. 500 BCE– 500 CE). In addition to brick architectural monuments, associated moats, and ponds, the Vat Komnou mortuary assemblage includes human burials, beads, ceramics, multiple pig skulls, and other faunal remains (Ikehara-Quebral 2010; Stark 2006). A total of 111 individuals were sorted and analyzed from 57 burial features excavated at the Vat Komnou cemetery by the Lower Mekong Archaeological Project (LOMAP) in 1999 and 2000. Participating LOMAP Institutions are: University of Hawai`i-Manoa (USA), Ministry of Culture and Fine Arts (Kingdom of Cambodia), Royal University of Fine Arts (Kingdom of Cambodia), University of Glasgow (Scotland, UK), and Scottish Universities Environmental Research Centre (Scotland, UK). There was extensive commingling of the burials at the Vat Komnou cemetery due, in part, to its apparent re-use through time (Ikehara-Quebral 2010). The Vat Komnou cemetery is one of the largest archaeological skeletal samples analyzed to date from Cambodia.

## Sample used in this study

- AB40: Male, 1890-1731 cal yBP (1885±30 BP, Poz-81120, date suspect due to C:N ratio of 3.75 – but reasonable based on archaeological context, and only slightly elevated C:N ratio)

Table S1: See attached file

**Table S2:** Outgroup f_3(Dinka; X, Y)

|  | Ami | Atayal | Borneo | Burmese | Cambodia_IA | Cambodian | CHB | Dai | Htin | KHV | Lahu | Lao | Miao |
| --- | --- | --- | --- | --- | --- | --- | --- | --- | --- | --- | --- | --- | --- |
| Ami |  | 0.2099 | 0.1951 | 0.1809 | 0.1674 | 0.1873 | 0.1963 | 0.1990 | 0.1903 | 0.1974 | 0.1946 | 0.1945 | 0.1970 |
| Atayal | 0.2099 |  | 0.1934 | 0.1796 | 0.1685 | 0.1859 | 0.1951 | 0.1981 | 0.1895 | 0.1961 | 0.1936 | 0.1929 | 0.1956 |
| Borneo | 0.1951 | 0.1934 |  | 0.1764 | 0.1691 | 0.1855 | 0.1866 | 0.1916 | 0.1901 | 0.1902 | 0.1890 | 0.1899 | 0.1886 |
| Burmese | 0.1809 | 0.1796 | 0.1764 |  | 0.1577 | 0.1739 | 0.1818 | 0.1810 | 0.1772 | 0.1801 | 0.1803 | 0.1782 | 0.1817 |
| Cambodia_IA | 0.1674 | 0.1685 | 0.1691 | 0.1577 |  | 0.1654 | 0.1668 | 0.1702 | 0.1681 | 0.1679 | 0.1682 | 0.1692 | 0.1654 |
| Cambodian | 0.1873 | 0.1859 | 0.1855 | 0.1739 | 0.1654 |  | 0.1837 | 0.1872 | 0.1858 | 0.1862 | 0.1852 | 0.1861 | 0.1850 |
| CHB | 0.1963 | 0.1951 | 0.1866 | 0.1818 | 0.1668 | 0.1837 |  | 0.1948 | 0.1869 | 0.1951 | 0.1934 | 0.1901 | 0.1972 |
| Dai | 0.1990 | 0.1981 | 0.1916 | 0.1810 | 0.1702 | 0.1872 | 0.1948 |  | 0.1919 | 0.1966 | 0.1950 | 0.1948 | 0.1964 |
| Htin | 0.1903 | 0.1895 | 0.1901 | 0.1772 | 0.1681 | 0.1858 | 0.1869 | 0.1919 |  | 0.1905 | 0.1905 | 0.1906 | 0.1894 |
| KHV | 0.1974 | 0.1961 | 0.1902 | 0.1801 | 0.1679 | 0.1862 | 0.1951 | 0.1966 | 0.1905 |  | 0.1934 | 0.1928 | 0.1951 |
| Lahu | 0.1946 | 0.1936 | 0.1890 | 0.1803 | 0.1682 | 0.1852 | 0.1934 | 0.1950 | 0.1905 | 0.1934 |  | 0.1914 | 0.1945 |
| Lao | 0.1945 | 0.1929 | 0.1899 | 0.1782 | 0.1692 | 0.1861 | 0.1901 | 0.1948 | 0.1906 | 0.1928 | 0.1914 |  | 0.1920 |
| Miao | 0.1970 | 0.1956 | 0.1886 | 0.1817 | 0.1654 | 0.1850 | 0.1972 | 0.1964 | 0.1894 | 0.1951 | 0.1945 | 0.1920 |  |
| Mlabri | 0.1894 | 0.1885 | 0.1911 | 0.1774 | 0.1662 | 0.1865 | 0.1856 | 0.1916 | 0.1950 | 0.1897 | 0.1900 | 0.1908 | 0.1880 |
| Myanmar_LNBA | 0.1840 | 0.1845 | 0.1806 | 0.1753 | 0.1622 | 0.1764 | 0.1887 | 0.1851 | 0.1794 | 0.1852 | 0.1846 | 0.1828 | 0.1867 |
| Naxi | 0.1905 | 0.1894 | 0.1829 | 0.1800 | 0.1624 | 0.1807 | 0.1944 | 0.1903 | 0.1843 | 0.1898 | 0.1912 | 0.1861 | 0.1931 |
| Nicobarese | 0.1875 | 0.1865 | 0.1912 | 0.1747 | 0.1692 | 0.1860 | 0.1822 | 0.1891 | 0.1912 | 0.1876 | 0.1874 | 0.1891 | 0.1847 |
| Onge | 0.1530 | 0.1527 | 0.1533 | 0.1493 | 0.1447 | 0.1519 | 0.1523 | 0.1531 | 0.1536 | 0.1527 | 0.1531 | 0.1532 | 0.1517 |
| Semende | 0.2021 | 0.1998 | 0.1925 | 0.1779 | 0.1680 | 0.1851 | 0.1907 | 0.1942 | 0.1894 | 0.1927 | 0.1909 | 0.1912 | 0.1923 |
| Thailand_BA | 0.1923 | 0.1893 | 0.1949 | 0.1744 | 0.2830 | 0.1838 | 0.1832 | 0.1824 | 0.1945 | 0.1818 | 0.1930 | 0.1893 | 0.1914 |
| Vietnam_BA | 0.1958 | 0.1949 | 0.1887 | 0.1771 | 0.1710 | 0.1848 | 0.1907 | 0.1940 | 0.1893 | 0.1930 | 0.1913 | 0.1911 | 0.1927 |
| Vietnam_NE | 0.1860 | 0.1866 | 0.1861 | 0.1727 | 0.1667 | 0.1810 | 0.1815 | 0.1881 | 0.1887 | 0.1863 | 0.1857 | 0.1863 | 0.1844 |

**Table S2 continued**
|  | Mlabri | Myanmar_LNBA | Naxi | Nicobarese | Onge | Semende | Thailand_BA | Vietnam_BA | Vietnam_NE |
| --- | --- | --- | --- | --- | --- | --- | --- | --- | --- |
| Ami | 0.1894 | 0.1840 | 0.1905 | 0.1875 | 0.1530 | 0.2021 | 0.1923 | 0.1958 | 0.1860 |
| Atayal | 0.1885 | 0.1845 | 0.1894 | 0.1865 | 0.1527 | 0.1998 | 0.1893 | 0.1949 | 0.1866 |
| Borneo | 0.1911 | 0.1806 | 0.1829 | 0.1912 | 0.1533 | 0.1925 | 0.1949 | 0.1887 | 0.1861 |
| Burmese | 0.1774 | 0.1753 | 0.1800 | 0.1747 | 0.1493 | 0.1779 | 0.1744 | 0.1771 | 0.1727 |
| Cambodia_IA | 0.1662 | 0.1622 | 0.1624 | 0.1692 | 0.1447 | 0.1680 | 0.2830 | 0.1710 | 0.1667 |
| Cambodian | 0.1865 | 0.1764 | 0.1807 | 0.1860 | 0.1519 | 0.1851 | 0.1838 | 0.1848 | 0.1810 |
| CHB | 0.1856 | 0.1887 | 0.1944 | 0.1822 | 0.1523 | 0.1907 | 0.1832 | 0.1907 | 0.1815 |
| Dai | 0.1916 | 0.1851 | 0.1903 | 0.1891 | 0.1531 | 0.1942 | 0.1824 | 0.1940 | 0.1881 |
| Htin | 0.1950 | 0.1794 | 0.1843 | 0.1912 | 0.1536 | 0.1894 | 0.1945 | 0.1893 | 0.1887 |
| KHV | 0.1897 | 0.1852 | 0.1898 | 0.1876 | 0.1527 | 0.1927 | 0.1818 | 0.1930 | 0.1863 |
| Lahu | 0.1900 | 0.1846 | 0.1912 | 0.1874 | 0.1531 | 0.1909 | 0.1930 | 0.1913 | 0.1857 |
| Lao | 0.1908 | 0.1828 | 0.1861 | 0.1891 | 0.1532 | 0.1912 | 0.1893 | 0.1911 | 0.1863 |
| Miao | 0.1880 | 0.1867 | 0.1931 | 0.1847 | 0.1517 | 0.1923 | 0.1914 | 0.1927 | 0.1844 |
| Mlabri |  | 0.1799 | 0.1834 | 0.1941 | 0.1538 | 0.1896 | 0.1792 | 0.1900 | 0.1893 |
| Myanmar_LNBA | 0.1799 |  | 0.1879 | 0.1787 | 0.1509 | 0.1810 | 0.2424 | 0.1967 | 0.1925 |
| Naxi | 0.1834 | 0.1879 |  | 0.1795 | 0.1518 | 0.1859 | 0.1878 | 0.1860 | 0.1792 |
| Nicobarese | 0.1941 | 0.1787 | 0.1795 |  | 0.1536 | 0.1886 | 0.1862 | 0.1867 | 0.1893 |
| Onge | 0.1538 | 0.1509 | 0.1518 | 0.1536 |  | 0.1534 | 0.1736 | 0.1485 | 0.1506 |
| Semende | 0.1896 | 0.1810 | 0.1859 | 0.1886 | 0.1534 |  | 0.1918 | 0.1915 | 0.1854 |
| Thailand_BA | 0.1792 | 0.2424 | 0.1878 | 0.1862 | 0.1736 | 0.1918 |  | 0.1852 | 0.1657 |
| Vietnam_BA | 0.1900 | 0.1967 | 0.1860 | 0.1867 | 0.1485 | 0.1915 | 0.1852 |  | 0.2059 |
| Vietnam_NE | 0.1893 | 0.1925 | 0.1792 | 0.1893 | 0.1506 | 0.1854 | 0.1657 | 0.2059 |  |

**Figure S1.**
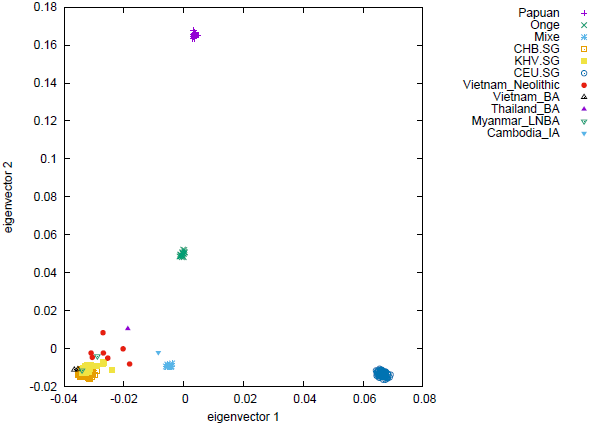
PCA with diverse non-Africans.

**Figure S2.**
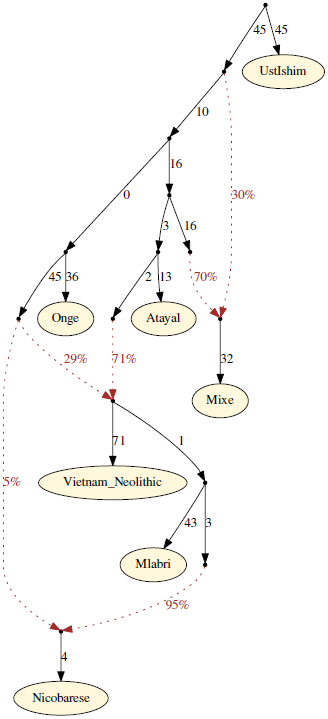
Basic admixture graph for Man Bac with present-day Austroasiatic-speaking populations. All *f*-statistics relating the populations are predicted to within 1.3 standard errors of their observed values. Omitting VN29, the two fitted mixture proportions are 28% and 6%, and all *f*-statistics are predicted to within 1.5 standard errors; omitting VN29 and VN40, the figures are 27%, 6%, and 1.7. Dotted lines denote admixture events, with proportions as shown. Branch lengths are given in units of 1000 times *f_2_* drift distance (rounded to the nearest integer).

**Figure S3.**
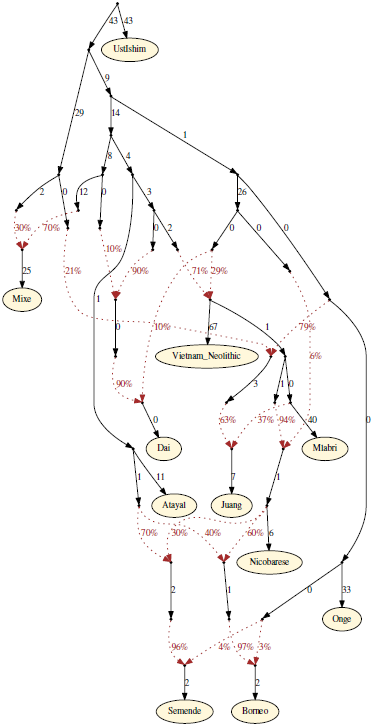
Extended admixture graph including Man Bac, present-day Austroasiatic-speaking populations, Dai, Borneo, Semende, and Juang. All *f*-statistics relating the populations are predicted to within 1.9 standard errors of their observed values. Dotted lines denote admixture events, with proportions as shown. Branch lengths are given in units of 1000 times *f*_2_ drift distance (rounded to the nearest integer).

